# Mapping neural activity during naturalistic visual and memory search

**DOI:** 10.1101/2025.07.27.667084

**Authors:** Joaquin E Gonzalez, Markus Bauer, Anthony J Ries, Juan E Kamienkowski, Matias J Ison

## Abstract

In everyday life, individuals often search for one of several items stored in memory. This cognitive process, known as hybrid search, is critical for tasks like navigating using landmarks. While the behavioral aspects of hybrid search have been extensively studied, the underlying neural mechanisms remain less understood. In this study, we combined concurrent magnetoencephalography (MEG) and eye movement recordings to investigate the oscillatory and evoked neural dynamics supporting hybrid search in naturalistic settings. Twenty-one participants (12 males, 9 females) performed a free-viewing task involving visual search for targets embedded in memory (hybrid search) across naturalistic scenes. Time-Frequency analyses revealed specific neural signatures during memory encoding, retention, and visual search. During encoding and retention, posterior alpha-band power decreased with memory load, reflecting heightened perceptual and mnemonic demands. During visual search, frontoparietal beta-band activity scaled with memory load, suggesting increased cognitive control. By aligning MEG signals to eye movement events and applying source reconstruction, we identified an early visually evoked lambda response, localized to V1, followed by a distributed P3m component, with maximum activation in the right inferior parietal lobe, that discriminated target from distractor fixations. Together, these findings demonstrate how oscillatory and evoked responses dynamically support hybrid search in naturalistic settings, revealing how memory, attention, and visual processing interact during active vision.

**Significance statement:** Understanding how the brain supports everyday tasks, like searching for familiar objects, requires studying cognition under realistic conditions, yet most knowledge comes from highly controlled lab tasks with simple stimuli. Here, we combined MEG and eye-tracking to examine how the brain supports free-viewing hybrid search -looking for any of several remembered items-across natural scenes. We found memory load modulations in posterior alpha oscillations during encoding and retention, as well as in frontoparietal beta oscillations during search. By time-locking neural signals to eye movements, we identified a robust marker of target detection. These findings provide direct evidence of how attention, memory, and vision interact in realistic settings, helping bridge the gap between controlled experimental paradigms and real-world cognition.

## Introduction

A fundamental premise in cognitive neuroscience is that using simple stimuli -designed to minimize confounds-, offers the “building blocks” for understanding complex real-world behavior. This approach has yielded insights across domains including attention and decision-making (Buschman & Miller, 2007; Gold & Shadlen, 2007; Posner & Petersen, 1990), yet little empirical evidence exists about how these building blocks interact.

In visual search, experimental paradigms often rely on artificial stimuli on uniform backgrounds. Neural studies of attention commonly restrict eye movements -using brief presentations (Desimone & Duncan, 1995) or requiring covert search (Çukur et al., 2013) to avoid M-EEG signal artifacts (Carl et al., 2012; Muthukumaraswamy, 2013). However, this overlooks the ubiquity of eye movements in natural vision. While controlled visual search studies (Treisman & Gelade, 1980) have provided key insights, visual search under real-world conditions remains largely unexplored (Peelen & Kastner, 2014). Recent studies combining eye-tracking with neural recordings have started to identify brain signatures of cognitive processes across naturalistic settings, such as recognizing faces in crowds (Kaunitz et al., 2014), virtual navigation (Stankov et al., 2021) watching movies (Nentwich et al., 2023). Building on these advances, we focus here on hybrid search, a common and ecologically relevant form of search in which individuals look for any of several memorized targets within visual environments (Schneider & Shiffrin, 1977; Wolfe, 2012; Barbosa et al., 2024). Hybrid search involves memory encoding, when potential targets are learned, and active visual search, where perceptual input is matched against stored representations.

Neuronal oscillations have been widely associated with attention, memory, and perceptual decision-making (Bauer et al., 2014; Fries, 2005; Rohenkohl & Nobre, 2011). In working memory, oscillatory activity recorded via M-EEG and intracranial methods shows frequency-specific links to memory representations (Axmacher et al., 2010; Roux & Uhlhaas, 2014). While early studies focused on theta and gamma rhythms (Jensen & Lisman, 1998), increasing evidence implicates alpha and beta oscillations in memory formation and maintenance (Hanslmayr & Staudigl, 2014; Staudigl et al., 2017). Alpha-band activity typically varies with memory load. Increases in posterior alpha have been linked to the suppression of irrelevant information (Jensen & Mazaheri, 2010; Klimesch, 2012), while decreases have been observed when perceptual detail must be actively maintained (Proskovec et al., 2019; van Ede, 2018; van Ede et al., 2017). Successful memory formation is also associated with changes in oscillatory amplitude or synchrony during encoding (Hanslmayr & Staudigl, 2014).

Invasive primate recordings during free-viewing show transient beta increases in V1 post-fixation (Maldonado et al., 2008; Ito et al., 2011). Beta oscillations have been linked to communication between dorsolateral prefrontal cortex and superior colliculus during eye movement tasks (Chan et al., 2015), and to decision-making in frontoparietal circuits during visual search (Pesaran et al., 2008; Stoll et al., 2016). Target detection is often marked by the P3-P3m component (Polich, 2007), a response observed in tasks ranging from oddball detection to free-viewing search (Kamienkowski et al., 2012).

Together, these findings suggest that oscillatory dynamics and evoked responses reflect core cognitive mechanisms underlying hybrid search. As most evidence comes from simplified paradigms, the “building blocks” identified in such settings remain disconnected from how they operate during complex behavior. Here, we combined MEG with eye-tracking to investigate fixation-related activity and task-related oscillations during a free-viewing hybrid search task. We tested four hypotheses: H1) Given that naturalistic stimuli require high perceptual detail, posterior alpha oscillations will attenuate with increasing memory load; H2) Alpha/beta power and phase during encoding will predict memory performance; H3) Because hybrid search requires maintaining multiple target representations, we predict that higher memory load will amplify beta-band activity as a marker of cognitive control; H4) Fixations on targets will elicit a P3m component, generalizing classic electrophysiological signatures to naturalistic contexts. By investigating how oscillatory and evoked responses support hybrid search under ecologically valid conditions, we aim to clarify how neural mechanisms -cognitive “building blocks”-operate during complex behavior, offering insights toward ecologically valid models of cognition.

## Methods

### Data acquisition

#### Participants

Twenty-one participants (12 males, 9 females) between the ages of 19 and 45 years (mean: 28.6 +/- 6.4) participated in this study. Participants were given an inconvenience allowance for their participation and provided written consent form after being instructed on every aspect of the experiment. The study was approved by the University of Nottingham School of Psychology Ethics Panel (ethics approval: F1349).

#### Experimental task

The experiment consisted of an all-new mapping (i.e. target changes every trial) hybrid search paradigm of 7 blocks of 30 trials each with an average duration of approximately 54 minutes. The stimuli were presented on a projected screen of approximately 38 cm width. Each trial was composed of a fixation dot of 750 ms, followed by a memorization/encoding screen. During this screen, the participants were presented with 1, 2 or 4 items corresponding to the memory set size (MSS), and the duration was 2, 3.5 or 5 seconds, respectively. During this time, participants were asked to memorize the items by freely exploring the screen. This was followed by a retention period of 1 second presenting a fixation dot, and finally, a search screen consisting of an image on which 16 items were embedded (Fig. 1). On half of the trials (“target present”), one of these items had previously been presented on the encoding screen; on the other half (“target absent”), none of the items in the search screen had been present during memorization. The search screen had a maximum duration of 10 seconds; during this time, participants could freely explore the screen and respond if any of the presented items during the encoding period were present or absent by pressing a button with the right/left thumb on a button pad. Participants could respond at any time during the search, which would end the trial. After each trial, there was a 1-second pause to rest. Participants had the choice to take a break between blocks.

**Figure 1.**
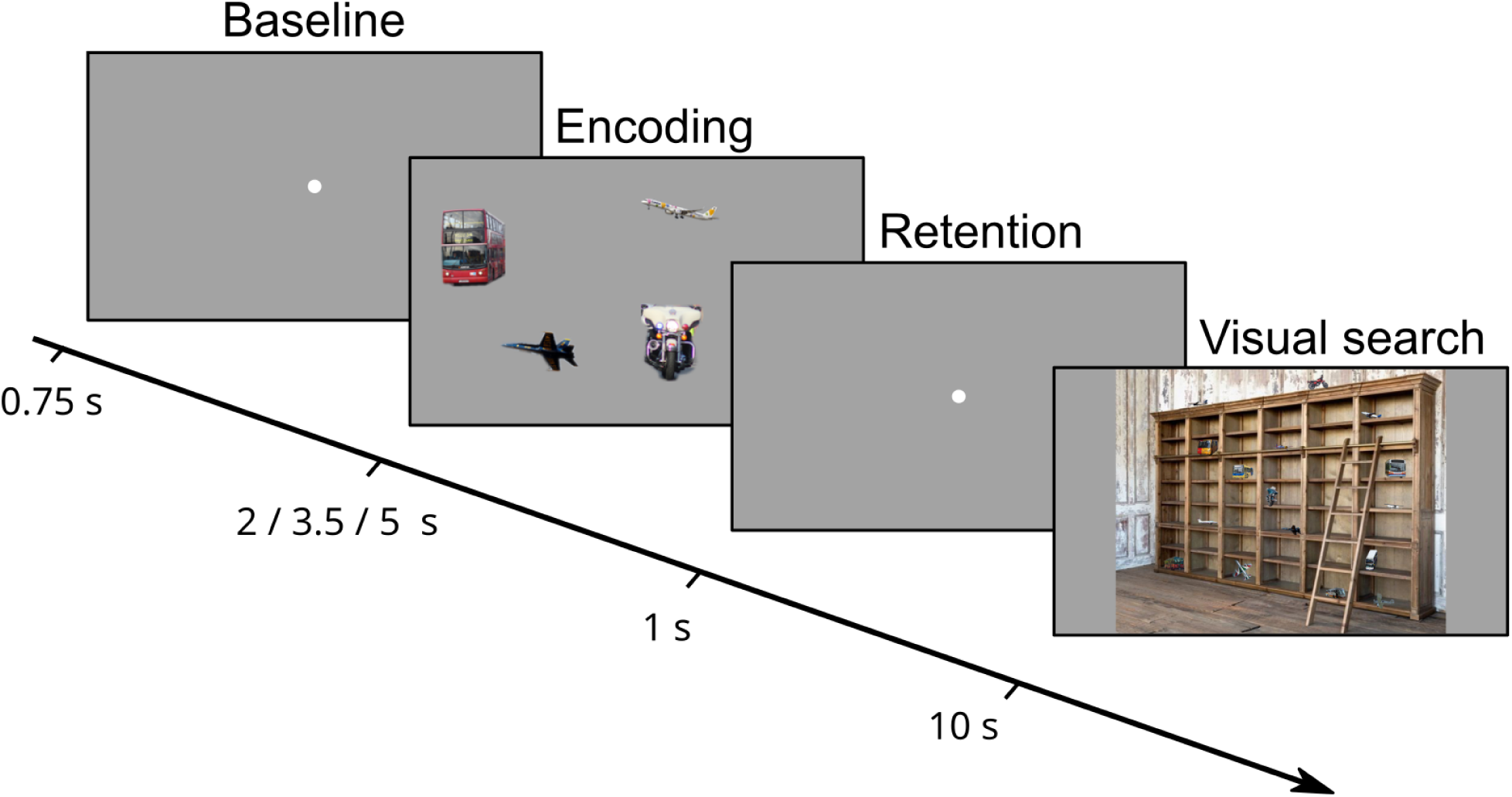
Overview of the experimental task. Each trial consisted of four sequential screens: An initial fixation cross, a memory encoding screen (items enlarged here for clarity), a retention fixation cross, and a naturalistic scene for free-viewing visual search.

In each search image (1280 × 1024 pixels), 16 individual items (objects, animals or full-body humans) were superimposed on a real-world background image (depicting outdoor scenes -e.g. forest-, and indoor scenes -e.g. shelf-) (Fig. 1). Every item and background image were taken from COCO (Lin et al., 2014) or ImageNet (Deng et al., 2009) datasets and appeared only one time across all images. A total of 210 images were presented. A bounding box was defined around each item, with a maximum size of 80×80 pixels. For a full characterization of the experimental stimuli see (Ruarte et al., 2025).

#### Head Digitization

Before the MEG acquisition, three electromagnetic coils were placed at three fiducial points on the participant’s head (nasion, left and right pre-auricular points), and digitization of these fiducial points and the head shape of the participant was carried out using a 3-D digitizer camera mounted on an iPad (Scanner app). During MEG data acquisition, the location of the participant’s head within the MEG helmet was measured by energizing these coils. The location of the MEG sensors was co-registered to the brain anatomy by matching the digitized head surface to the head surface extracted from the anatomical MR image following (Brookes et al., 2011).

#### MEG recording

During the experiment, brain activity was recorded using a CTF 275 MEG, inside a magnetically shielded room. The experiment was presented to the participants through a projector, projecting an image onto a screen in the magnetically shielded room. On each trial, triggers were sent from the experiment display PC to the eye-tracker and MEG, indicating the beginning of the trial and the beginning and end of the search screen. The CTF system was configured to record at 1200 Hz and to apply 3rd order spatial gradient correction on the data to minimize the influence of environmental magnetic noise. MEG recordings of the empty room were performed regularly to use as background noise for source estimation.

#### ET recording

A MEG-compatible Eye tracker (ET) (Eyelink 1000 Plus, SR Research, Ontario, Canada) was used to record eye movements at a sampling rate of 1,000 Hz. The eye tracker was set to monocular mode to record the participant’s dominant eye and placed beneath the projector screen at approximately 68 cm from the participants’ eyes. The eye-tracker 9pt calibration took place at the beginning of the first block and was recalibrated between blocks when necessary. Calibration errors were kept below 0.5 deg throughout the experiment. The ET data was stored in the Eyelink host PC, but it was also converted to an analogue signal using Eyelink’s Digital to Analogue converter and stored as extra analogue channels with the MEG data.

#### MRI Acquisition

For 14 of the 21 participants, a structural MRI was collected using a standard sMPRAGE T1 sequence in a 3T Siemens scanner. The scanning parameters were as follows: TR = 2,000 ms, TE = 2.01 ms, TI = 880 ms, flip angle = 8 degrees, FOV = 256 × 256 × 160 mm, 1 mm isotropic voxel. For the remaining participants, the MNI 305 template brain was used in the later source modeling analysis.

### Pre-processing

#### ET signal pre-processing

The ET data was taken from the analogue channels on the MEG to facilitate the synchronization with the MEG gradiometers’ data. The ET data was first rescaled from Volts to pixels following the Eyelink manual, then blinks were detected and removed by defining them as missing values in the signal, and then saccades and fixations longer than 50 ms were detected using Remodnav (Dar et al., 2021), an implementation of the Nystrom and Holmqvist’s velocity-based eye movement detection algorithm (Nyström & Holmqvist, 2010) (see Supplemental Texts S1 for details on the parameters used for the detection). Fixations on encoding and visual search screens were classified as fixations on targets, distractors, or ‘none’, depending on their distance to the items center using a 70 pixel (1.13 deg) radius threshold.

Finally, to determine if a delay was introduced by the Digital-Analogue Converter (DAC), epochs were time-locked to saccade onset, and the evoked response aligned to this onset was computed for the Grand Average of participants (GA). The evoked response presented a first peak approximately 10 ms before the onset of the saccades. This peak is known to take place right after saccade onset (Carl et al., 2012), so a -10 ms shift was introduced in the ET data to compensate by removing the first 12 samples (sampling rate = 1200 Hz) and padding the last 12 samples with empty values. After determining the DAC delay and compensating for it, the detection of fixations and saccades was re-run with the same parameters to correct the timing of the ocular events.

#### MEG signal pre-processing

The processing and analysis in this work were performed using MNE-python [MNE]. First, a 1 Hz bandwidth notch filter was applied to filter line noise at 50 Hz, 100 Hz, and 150 Hz. Bad channels were manually selected based on visual inspection for each participant, for an average of 2.3 +/- 3.3 bad channels per participant. These channels were Interpolated from neighboring channels using minimum-norm estimation. Afterwards, noisy intervals on the signal were identified and labeled as “bad”, by visual inspection, and muscular artifacts were labeled using an automatic method from MNE package v. 1.5.1 (*annotate_muscle_zscore*) on high frequency filtered data (110 - 140 Hz). These intervals were excluded from all the analysis on this work.

An Independent Component Analysis (ICA) was applied to every subject separately to remove ocular, muscular, cardiac, and noisy components. To this end, a copy of the preprocessed MEG signal was first downsampled at 200 Hz and filtered between 1 and 40 Hz, using 64 ICA components. The spectrum, topography, epochs (-50 to 100 ms) and evoked signal around saccades with left direction were plotted for each component individually and analyzed to determine if that component contained electrical noise, cardiac artifacts, and ocular artifacts. Since eye movements are present throughout the experiment, we followed the procedure introduced in (Plöchl et al., 2012) to identify eye-related components. In brief, this approach compares the variance around saccades with the variance around fixations and sets a semi-automatic threshold of 1.1 (higher variance associated with saccades) to define eye-related components. Once the artifactual components for each subject were defined (average 11.6), they were removed from the original unfiltered data and saved.

After ET signal processing and synchronization, events were marked on the MEG data, indicating each screen onset, and fixations, saccades, and button responses, to epoch the MEG data around times and conditions of interest.

#### Source estimation

Source estimates were computed using LCMV beamformer spatial filters following the procedure from (Pan et al., 2023). To this end, first, the head model of each participant was defined using its individual anatomical MRI, co-registered with the MEG data through the fiducial markers from the head digitization during MEG recording preparations. The MRI data was processed and segmented using FreeSurfer to extract the Boundary Element Method (BEM) by assigning each voxel of the MRI data to a tissue class. For the 7 participants lacking an MRI scan, the MNI 305 template was used as the head model.

The BEM was later used to construct two source models for each participant: a volume model using a 5 mm-spaced isotropic grid and a surface model using ico-4 spacing of 2562 sources per hemisphere. These models were then used to define forward models for volume and surface estimates.

The LCMV filters were constructed using the covariance matrices of the empty room recordings, and the covariance matrices of the whole MEG recording for each participant. These matrices were subject to regularization of 0.05 (5% of the sensor power) before inverting them in the beamformer calculation. Using all the MEG recordings provides a robust estimation of the covariance matrix and also avoids the need for biased LCMV filters for each condition from which sources would be estimated. To construct spatial filters for specific frequency bands (theta: 4 - 8 Hz, alpha: 8 - 12 Hz, beta: 12 - 30Hz), the sensor data was first filtered with a 4th order Butterworth IIR filter, and source estimates were computed from the filtered sensor data (Barratt et al., 2018). The empty room recordings were subject to the same pre-processing and artifact removal methods as the MEG recording before being used in the LCMV filter. At each source location, dipole activity was estimated in three orthogonal directions using a vector beamformer. Then, the scalar amplitude was computed as the norm of that vector, representing the total source power independent of orientation. These LCMV filters were finally applied to epoched or evoked time series and covariance matrices of epoched data. The results in individual participants’ volume source space were then morphed onto a common space, in this case, the MNI 305 template, where homologous points across participants were located at the same location, allowing for the signals and power estimates in source space to be directly averaged across participants.

Surface source estimates were divided into 148 discrete cortical regions, defined based on the Destrieux atlas (Destrieux et al., 2010). Singular value decomposition was applied to the time courses within each label, and the first right-singular vector was taken as the representative label time course. This yielded one signal per cortical region, from which time-frequency power was computed as detailed below.

### MEG analysis

#### Time-frequency analysis

The time-frequency analysis for real or virtual sensors was performed using Morlet wavelet filters in 1 Hz bands with MNE (Gramfort et al., 2013). The Morlet wavelets used for each frequency had the number of cycles defined as frequency / 2, which balanced the temporal and spectral resolutions.

#### Connectivity analysis

To investigate the amplitude coupling between different brain regions, a connectivity analysis was performed using amplitude envelope correlation (AEC) (Boto et al., 2021). Surface sources were estimated from the epoched MEG data of each participant corresponding to the fixations to the target and distractors using an LCMV Beamformer filter and aggregated into regions (following the procedure described above). The envelope of the signal for each region was extracted, and the correlation between each pair of time-series was computed, using pairwise orthogonalization for leakage correction (Boto et al., 2021; Rier et al., 2024). This method is robust against spurious synchronizations due to spatial leakage (Sareen et al., 2021). The resulting adjacency matrix was standardized and then averaged across participants when presenting grand average results. For comparing the connectivity of two conditions, the connectivity matrices of each participant were subtracted without any normalization or standardization, obtaining one difference connectivity matrix per participant. That matrix was then standardized by estimating z-scores for each participant and averaged across participants to get the difference in connectivity for the average of subjects.

To represent the connectivity of brain regions we use three visualization methods: Connectome plots, that show nodes located in the brain, and the links between those nodes, brain surface plots showing the connectivity degree of each region, and the connectivity matrix itself. The connectome plots present the strongest connections in the connectivity matrix. This is achieved by setting a threshold as the 150th highest absolute value of the matrix. Links with strictly higher absolute values than that threshold are kept, resulting in 149 links at most depending on repeated values in the matrix. The connectivity degree plot shows the sum of the connectivity weights of each node or region (connectivity degree for continuous valued networks).

To assess the significance of the difference in connectivity between hemispheres, the average connectivity matrix was used to select the 150 highest links in absolute value. Those links were extracted from the connectivity matrix of each subject and split into left and right hemispheres and used in the analysis. Connectivity values were averaged for each participant and hemisphere separately, yielding a left and right connectivity index for each participant. Finally, the left and right connectivity values were subject to a Wilcoxon signed-rank test.

#### Cluster statistics

The statistical significance of the results (both in time-frequency space or power source space and virtual sensor signals) was assessed by nonparametric cluster-based permutation tests implemented in MNE-python (Gramfort et al., 2013; Maris & Oostenveld, 2007). For time-frequency sensor-space data, the clustering was estimated on 3 dimensions (time x frequency x sensors) on the basis of spatial, spectral, and temporal adjacency. For source-space time series data, clustering was based on both spatial and temporal adjacency. Finally, for average power data in source-space, clusters were computed considering only spatial adjacency.

Mass-univariate comparisons were thresholded using a t-value of 2.84 (corresponding to p=0.01, d.f. = 20) to define candidate clusters. For each subject, the difference between conditions (task vs. baseline, or high vs. low memory load) was computed individually, and submitted to a one-sample permutation test, with 1024 permutations. Clusters with permutation-based p-values smaller than 0.05 were considered statistically significant.

## Results

### Behavior and Eye Movements

Participants were asked to memorize a set of items of different sizes (MSS = 1, 2, 4) and then search for any of them in a natural image including several items (Figure 2A). Response times showed a characteristic logarithmic increase with memory set size (Figure 2E) (Barbosa et al., 2024). Eye movements analysis showed typical distributions for fixation durations, saccade amplitude, angular distribution, and main sequence (Figures 2 C-D, F-G), consistent with established patterns in eye tracking data across tasks (Otero-Millan et al., 2008).

**Figure 2.**
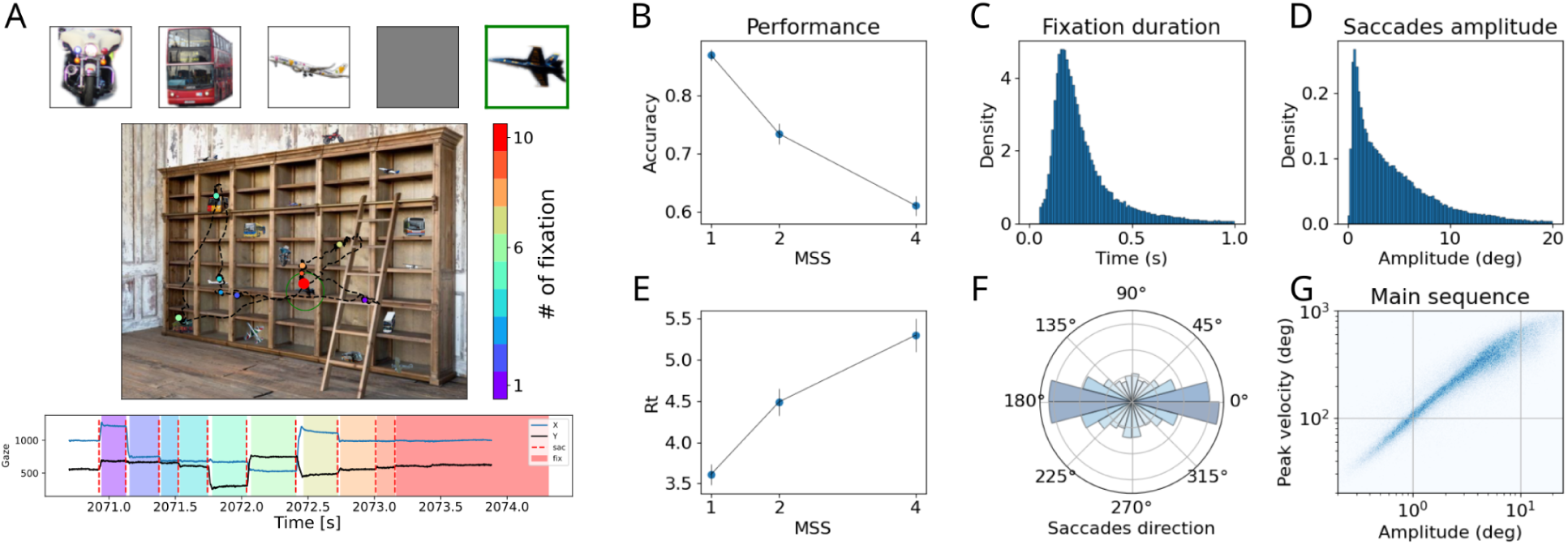
Behavioral performance and eye movement statistics during visual search. **(A)** Example trial for a memory set size MSS of 4. The top panel shows the memory set; the middle panel presents the scanpath over the visual search (VS) display with saccades as dotted black lines and fixations as colored dots indicating their temporal order (colorbar); the bottom panel plots horizontal (X) and vertical (Y) gaze positions over time. **(B)** Mean accuracy (+-S.E.M.) across different memory set sizes. **(C)** Distribution of fixation durations during visual search. **(D)** Saccade amplitude distribution. **(E)** Mean reaction times (+-S.E.M.) across different memory set sizes. **(F)** Saccade angular distribution. **(G)** Main sequence plot showing the relationship between saccade amplitude and peak velocity during visual search.

### Task-related brain activity

Figure 3 shows an overview of the power changes over the parietal and occipital sensors throughout the task across all memory loads. In the encoding phase, oscillatory power increased in the theta band (4-8 Hz) approximately 100 to 200 ms after screen onset and then decreased strongly around 500 ms in the alpha (8-12 Hz) and beta band (12-30 Hz). In the retention interval, alpha power significantly decreased with memory load, both in sensor space and in source space.

**Figure 3.**
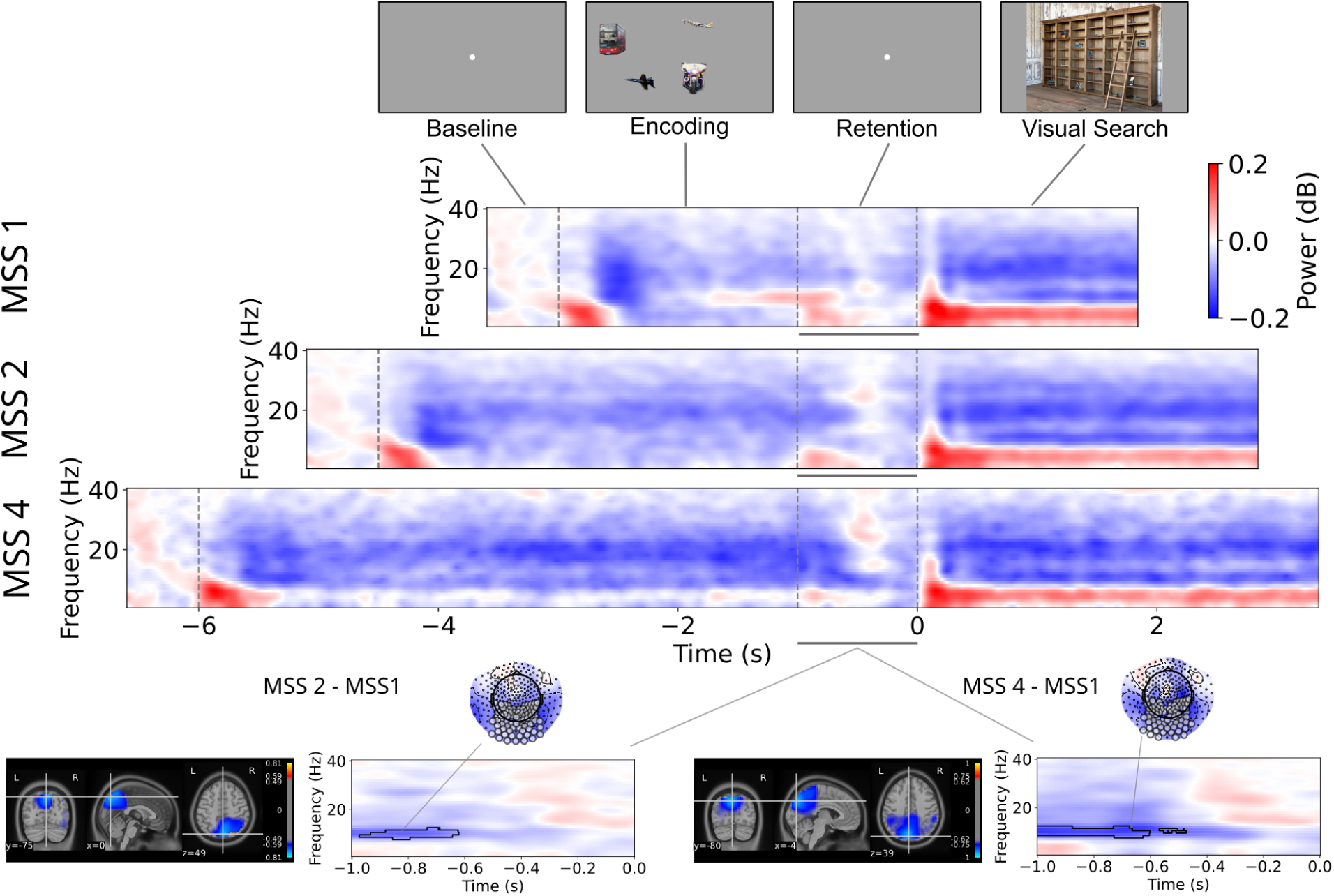
Time-frequency spectrograms (TFS) and memory load effects. Schematic of the task (top panel) and TFSs across memory set sizes (middle panels) from parietal and occipital sensors. Difference in spectral power (bottom panels) for medium/high vs. low memory set size during the retention period, shown in source space (bottom left) and sensor space (bottom right). Significant clusters, delimited by black contour lines, were estimated with a permutations test (see Methods). These clusters reflect time-frequency points where at least half of the sensors of the region were significant (p < 0.05). Topographic plots show the power distribution with grey circles marking significant sensors. Source space voxels with significant power differences between memory set sizes show maximum differences in the superior parietal lobe (Brodmann 7) in both cases.

The bottom panel of Figure 3 shows significant differences in alpha power between medium (MSS=2) and low (MSS=1) memory load conditions, as well as between high (MSS=4) and low (MSS=1) conditions. In contrast, there were no significant differences between high and medium memory load conditions. These results support alpha power suppression in posterior regions as a reliable marker of WM load, with WM reaching its maximum capacity for encoding natural objects at medium memory load. Source-level power during encoding and retention periods was estimated using an LCMV Beamformer applied to covariance matrices derived from the corresponding epochs, filtered in the alpha band (see Methods Section). To study the effect of memory load during the encoding and retention periods in source space, the difference between the source power for each load condition was computed, and a permutations test was performed on source space power estimates. This resulted in significant differences between high vs low memory load (p = 0.0029) and med vs. low memory load (p = 0.0039) conditions (Fig. 3, bottom panels). In both cases, the largest significant differences were found in the superior parietal lobe (Brodmann 7). No significant differences were found for high vs. medium memory load conditions. The decrease in alpha power with memory load was not specific to the retention period. During memory encoding, we found significant differences in the alpha parietal power between high (MSS=4) and medium (MSS=2) when compared to low (MSS=1) memory load conditions (see Fig. S1).

We also investigated whether differences in amplitude or synchronization of oscillations were predictive of successful memory performance. However, no significant differences in power nor phase synchronization (estimated using inter-trial phase coherence) were found when comparing correct and incorrect trials (Fig. S2). This analysis was performed both on a “whole trial” basis, with the overall power and phase during the encoding period and also aligning the data to individual saccades to items, since previous studies have reported stronger alignment of oscillations with saccade onset rather than fixation onset (Pan et al., 2023).

### Effect of memory load on fixation-related responses

To assess the effect of memory load on fixation-related responses in visual search, the time-frequency responses aligned to the onset of distractor fixations for each brain region were computed. The contrast between fixations under high (MSS=4) and low (MSS=1) memory load is presented in Figure 4, which shows two distinct clusters of regions with consistent modulation of responses with the memory load approximately 100 ms after fixation onset. Significant activations were most prominently observed in the beta band. The brain regions which presented such activations are shown on the left, extending from frontal to parieto-occipital regions.

**Figure 4.**
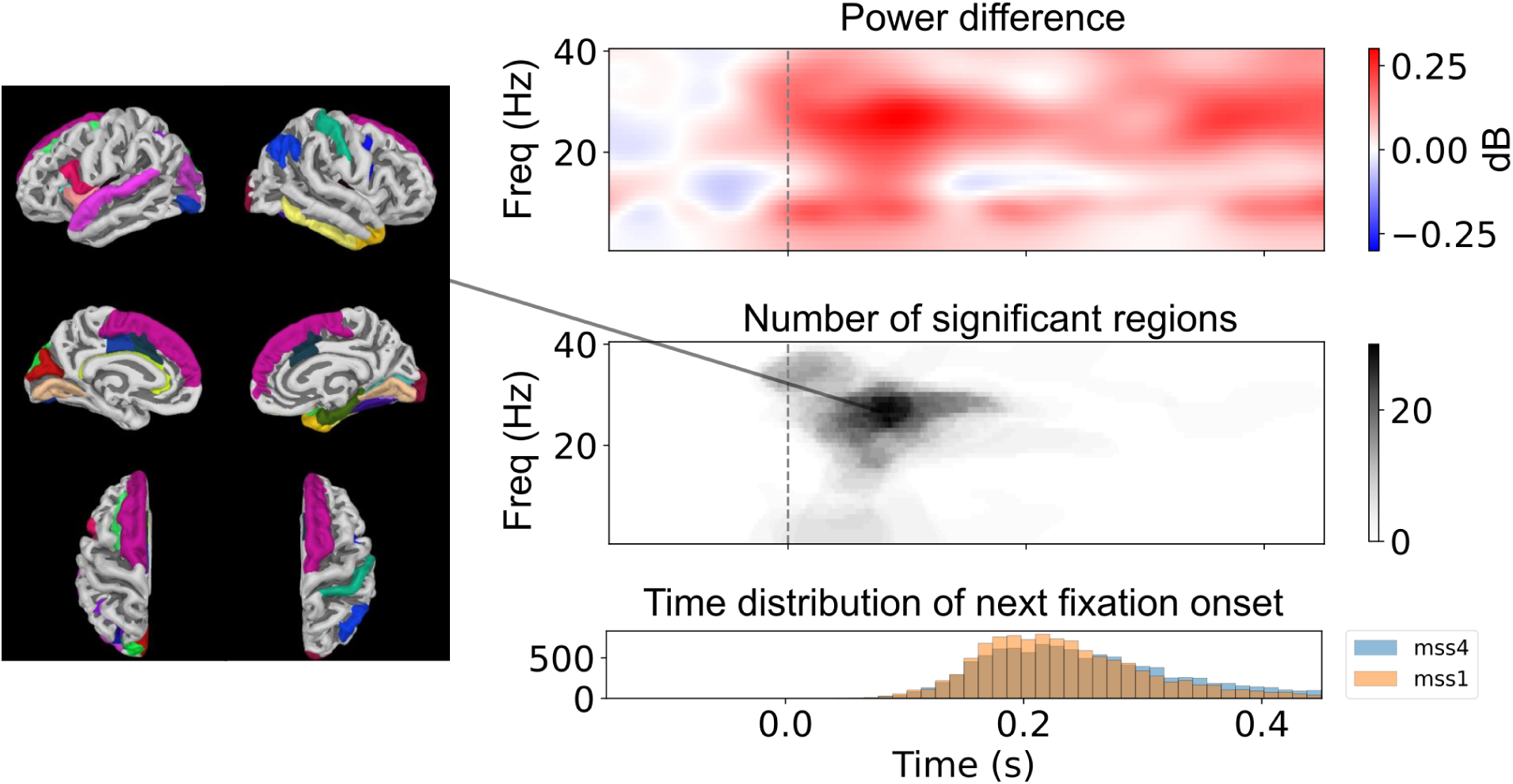
Modulation of fixation-related power by working memory load. Power differences in neural responses to distractor fixations during visual search are shown for trials with high (MSS=4) versus low (MSS=1) memory set sizes. The top panel illustrates time-frequency representations of power modulation, with two prominent clusters (left panels) indicating significant differences across a large number of regions. The bottom panel shows the distribution of the next fixation onset (i.e., current fixation duration + next saccade duration) for both memory load conditions, confirming that observed differences arise during the current fixation.

We repeated this analysis, aligning epochs to saccade onset (Pan et al., 2023). We obtained similar results using this alternative approach, both in terms of the number of regions presenting significant differences, as well as the effect of beta band oscillations on memory load (see Supplemental Fig. S3).

### Target recognition

To analyse the effect of target recognition, fixation-related fields for fixations on targets and distractors in the visual search phase were computed. These analyses included only fixations (for distractors and targets) from trials where the target was present, and responses were correct.

Due to the imbalance between conditions (around 15 fixations on distractors vs. 1 fixation on a target per search image), fixations to distractors were randomly subsampled for each participant. The evoked responses were computed for each sensor, participant and condition (target vs. distractor) separately and then averaged across participants to generate a Grand Average response for each channel and condition. Figure 5A shows the time courses of the fixation-related responses to targets and distractors in temporal sensors. Both target and distractor fixation-related responses present a prominent peak at approximately 100 ms, reflecting an occipital response to fixation onset, which has previously been referred to as the visually evoked lambda response in concurrent EEG and eye movements studies (Kaunitz et al., 2014; Ries et al., 2016, 2018). A statistically significant difference between targets and distractors, the target-related P3m, clearly emerges from ∼250 s to ∼350 ms after fixation onset on temporal sensors.

**Figure 5.**
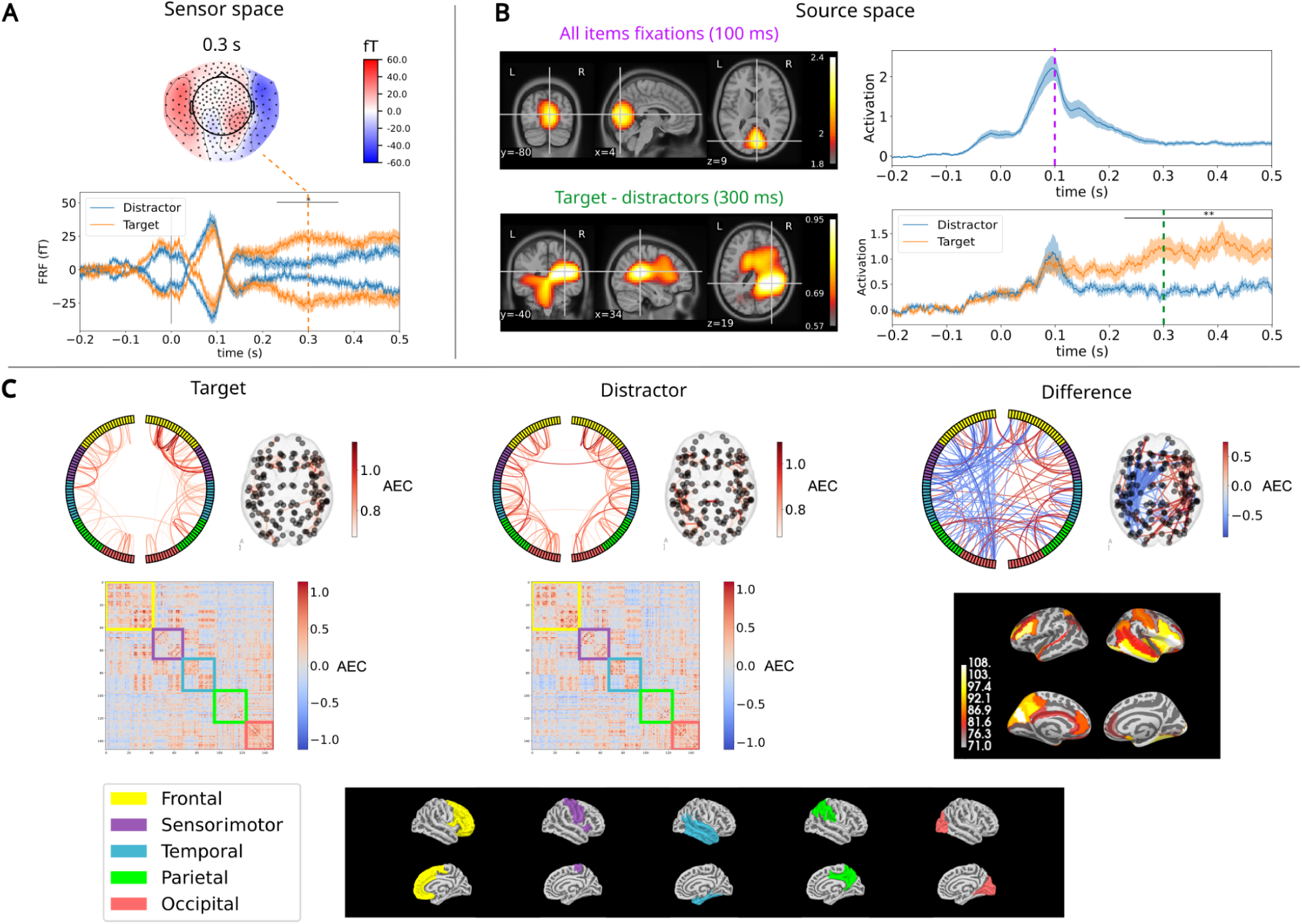
Evoked fixation-related responses to targets and distractors during visual search. **(A)** Sensor-level time series and topographic maps. Time series represent participant-averaged signals from lateralized temporal channels (highlighted in grey on the topographic maps), comparing fixations on targets (orange) and distractors (blue). A consistent peak occurs approximately 100 ms after fixation onset across both conditions. Target fixations show a second, significantly larger amplitude response between 250 and 350 ms (p=0.007 for right, p=0.01 for left electrodes). The topographic map shows the mean activation at 300 ms for target fixations. **(B)** Source-level estimates. The top panel shows activation at 100 ms localized to V1 (MNI: [4, -80, 9]). The bottom panel shows differential activation between target and subsampled distractor fixations at 300 ms, with a maximum activation in the right inferior parietal lobe (MNI: [34, -40, 19]). Virtual electrode time series from 0.125 cm3 regions centered on these specified voxels reveal significant differences between target and distractor fixations from approximately 250 ms to 500 ms (p=0.002 for V1, p=0.0005 for the tempo-parietal junction). (**C)** Connectome plots, depicted as circular graphs, connectivity matrices, and glass brains, during target and distractor fixations. Values are z-score standardized. The connectome plots show the strongest 150 links from the connectivity matrices. The connectivity degree plots (bottom right) show the sum of the connectivity weights involving each region.

Figure 5B shows, in the top panel, source activations for the fixation-related field at 100 ms after fixation onset, localized in the V1 region. The time course of the associated virtual sensor, taken from the location of maximum activation, is shown on the right. The bottom panel shows significant differences between target and distractor fixations from 250 ms onwards. The virtual electrode with maximum activation at 300 ms is present around the temporo-parietal junction and the broader right inferior parietal lobe (Brodmann areas 39 and 40), as shown in the activation map (bottom), although increased activation is visible across many areas. Statistical significance was assessed through a permutations test (p = 0.00097, Fig. S4).

To further characterize the widespread activation observed during target detection, we performed a functional connectivity analysis during target and distractor fixations. Figure 5C shows connectivity matrices and connectome plots for each condition, as well as their difference (target - distractors). Both target and distractor fixations show strong bilateral temporal-frontal synchronization. Consistent with the analysis of fixation-related fields (Fig 5B), the contrast between conditions shows a strong lateralization effect, with significantly higher connectivity for fixations on targets in the right hemisphere (Wilcoxon signed-rank test, p = *0.00000095*). Notably, all participants showed stronger connectivity for distractor fixations on the left hemisphere, and stronger connections for target fixations on the right hemisphere (Fig. S5).

## Discussion

In this work, we show the contribution of brain oscillations and fixation-related activity to hybrid visual and memory search tasks. Memory-dependent effects were observed in parietal alpha power, with attenuation starting in the encoding phase and persisting throughout the retention interval. We also found memory load effects on fixation-related responses, particularly in the beta band, across a large number of frontal and parieto-occipital regions. Neural signatures of target detection during the visual search phase emerged from fixation-related responses, as seen in sensor space, source space, and functional connectivity analyses. The timing and topography of these effects align with findings from fixed-gaze neuroimaging studies (Herrmann & Mecklinger, 2000; Kok, 2001; Polich, 2007), free-viewing EEG (Gordon et al., 2024; Kamienkowski et al., 2018; Kaunitz et al., 2014), as well as intracranial and simultaneous EEG-fMRI recordings (Bledowski et al., 2004; Halgren et al., 1998).

Consistent with earlier findings reporting alpha/beta desynchronization during encoding (Fukuda et al., 2015; Griffiths et al., 2019), we found significant markers of such effect aligned to stimulus presentation. This started during the encoding phase, in line with previous fixed gaze working memory studies (Proskovec et al., 2019) and persisted during the retention period (Chen et al., 2022). The attenuation of posterior alpha oscillations was modulated by memory load, thus supporting our experimental hypothesis H1. Our findings are consistent with a large corpus of works reporting either an amplification of alpha oscillations linked to distractor inhibition, where alpha amplifications in non-relevant task regions are thought to reflect the suppression of distracting stimuli and facilitate more efficient communication between task-relevant regions, or an attenuation of alpha oscillations in task-relevant regions, where alpha oscillations are thought to serve a genuine mnemonic function (Bauer et al., 2014; Erickson et al., 2019; Jensen & Mazaheri, 2010). Given the involvement of posterior regions during encoding and retention of fine-grained perceptual representations, as required by the task, our results support the role of posterior alpha oscillations and the maintenance of working memory representations (van Ede, 2018).

Contrary to our second hypothesis (H2), we found no differences in phase synchronization or activity when comparing oscillations linked to trials in which the target was found or not. As reviewed by (Hanslmayr & Staudigl, 2014), evidence from noninvasive M-EEG studies and intracranial EEG during the encoding of memories typically shows changes in amplitude or phase synchronization linked to successful memory formation. These changes manifest as increases or decreases in theta and alpha frequency bands, and as decreases in the beta band. They are also dependent on the way in which information is processed during encoding, with alpha and beta power decreases predicting memory formation only in semantic tasks (Hanslmayr & Staudigl, 2014). The underlying hypothesis is that successfully encoded memory traces are reinstated during retrieval. There are a number of possible reasons why we did not observe this in our experiment. First, we could only conduct this analysis for the high memory load condition, as there were fewer incorrect trials for the other set sizes. Second, while memory retrieval is usually assessed directly in memory experiments, in our experiment, successful trials corresponded to targets in which the target was found, thus leaving the possibility that the target was presented but not fixated on during the visual search process. Third, with the exception of (Staudigl et al., 2017), most of the existing evidence comes from experiments conducted under fixed gaze in which, arguably, differences between networks involved in successful and unsuccessful encoding might be easier to detect.

We observed effects of memory load on fixation-related responses, consistent with our hypothesis H3. When participants attempted to recognize a target among a series of memorized items, beta oscillations increased with memory load around 100 ms post-fixation in a large number of frontal and parietal regions. Frontal beta oscillations have traditionally been associated with top-down control mechanisms (Miller et al., 2018), task demand and attentional effort (Donner et al., 2007; Gross et al., 2004; Siegel et al., 2012; Stoll et al., 2016). During a free-viewing visual search task, Pesaran and colleagues found that beta coherence was stronger during free-choice compared to instructed decisions, suggesting that top-down signals from frontal regions shape decision dynamics (Pesaran et al., 2008). In another experiment in monkeys performing a test of cognitive control, Stoll and colleagues found a significant increase in beta oscillations only when cognitive control was required (Stoll et al., 2016). Our findings support that the beta modulation in temporal and frontal regions is related to early access to working memory, associated with the recognition attempt, which becomes more pronounced under higher memory demands.

In line with our hypothesis H4, we found robust neural markers of target detection. The neural mechanisms underlying target detection in visual search have been extensively studied using electrophysiological and neuroimaging approaches. One well-established neural signature of target detection is the P3 component (and its magnetic counterpart, P3m), which emerges around 300-600 ms after stimulus onset and has traditionally been linked to contextual processing of target-related information (Polich, 2007). The P3m topography we observed, with a maximum over temporal sensors, is consistent with previous fixed gaze works (Herrmann & Knight, 2001), and generalizes previous findings of concurrent EEG and eye tracking recordings during free viewing visual search (Kamienkowski et al., 2018; Kaunitz et al., 2014; Touryan et al., 2017). Our results show a widespread network involved in target processing. Intracranial recordings from neurosurgical patients have identified P3 generators in widespread regions, including the hippocampus, superior temporal sulcus, ventrolateral prefrontal cortex, and intraparietal sulcus (Halgren et al., 1998). By applying source reconstruction techniques for MEG recordings, we found maximum activation in the right inferior parietal lobe. (Bledowski et al., 2004) used a three-stimulus oddball task with combined EEG and fMRI recordings and reported activation of a widespread cortical network, with pronounced activation in parietal regions including the inferior parietal lobe and the posterior parietal cortex, as well as a strong lateralized effect with greater activation associated to targets in the right hemisphere. This is also consistent with the results of our functional connectivity analysis for fixations on targets and distractors, which revealed increased brain connectivity in the right hemisphere for targets, while the connectivity in the left hemisphere showed larger connectivity for fixations on distractors.

There are a number of limitations in our study. First, while traditional neuroimaging experiments normally present carefully designed isolated stimuli on blank backgrounds, we have digitally manipulated naturalistic scenes to add target and distractor stimuli. We believe that using naturalistic stimuli is advantageous to better approximate how the brain functions ‘in the wild’ (Sonkusare et al., 2019). However, these experiments lack control over specific task conditions. Second, unlike fixed-gaze experiments, our task encouraged participants to perform as many saccadic eye movements as needed. We built on extensive previous experience from concurrent M-EEG and eye movements recordings (Carl et al., 2012; Kaunitz et al., 2014; Plöchl et al., 2012) to identify and correct for artifactual components, but we cannot rule out the contamination of eye movements in the processed data. Moreover, even if the muscular artifacts are perfectly accounted for, the processing of information during each fixation in natural vision will likely be influenced by the activity associated with the previous fixation. There is good evidence to support the robustness of the fixation-related approach, including the identification of a robust lambda wave component. This component, estimated from co-registered MEG and MRI signals, localized to V1 matching the localization reported by a previous EEG study (Kazai & Yagi, 2003). Further analysis methods may be necessary to fully disentangle overlapping effects from consecutive fixations, and their source localization in isolation. Deconvolution methods (Care et al., 2023; Crosse et al., 2016; Lalor et al., 2006; Litvak et al., 2013) have been shown to robustly handle such unconstrained scenarios. Future work should implement these methods to further explore the neural mechanisms underlying visual search, focusing on fixation- and saccade-level representations, how these processes scale from short time windows to full-trial activity, and combine these methods with precise source localization from MEG.

In conclusion, our study sheds light on the neural mechanisms of visual search under naturalistic conditions, emphasizing the role of alpha and beta oscillations, and fixation-related responses in guiding target detection and working memory processes. By bridging the gap between traditional laboratory studies and real-world scenarios, our findings contribute to a deeper understanding of visual cognition and its neural underpinnings.

## Code availability

The code for the preparation of the data and the analysis is available in https://github.com/jegonza66/MEGEYEHS

## Supplemental Material

https://docs.google.com/document/d/1-PlerRgyx_Oj0dGcRTQAjmTuDWlSokCZ_66NbPr-QFU/edit?usp=sharing

