## Supplementary information for "Mapping neural activity during naturalistic visual and memory search"

### Texts S1: Eye Tracker Data

REMoDNaV (Dar et al., 2021), with the implementation found here: https://github.com/psychoinformatics-de/remodnav was used to detect eye movement events.

Parameters used for the detection:
--savgol-length: 0.0195
--min-pursuit-duration: 2 (To avoid getting smooth pursuit events)
--max-pso-duration: 0.0 (To avoid having Post saccadic oscillation events)
--min-fixation-duration: 0.05 (Min fixation duration 50 ms)
--max-vel: 5000

Additionally, saccades with peak velocity over 1500°/s and fixations with an amplitude over 1.5° were removed.

### Figure S1: Memorization Time-Frequency Response

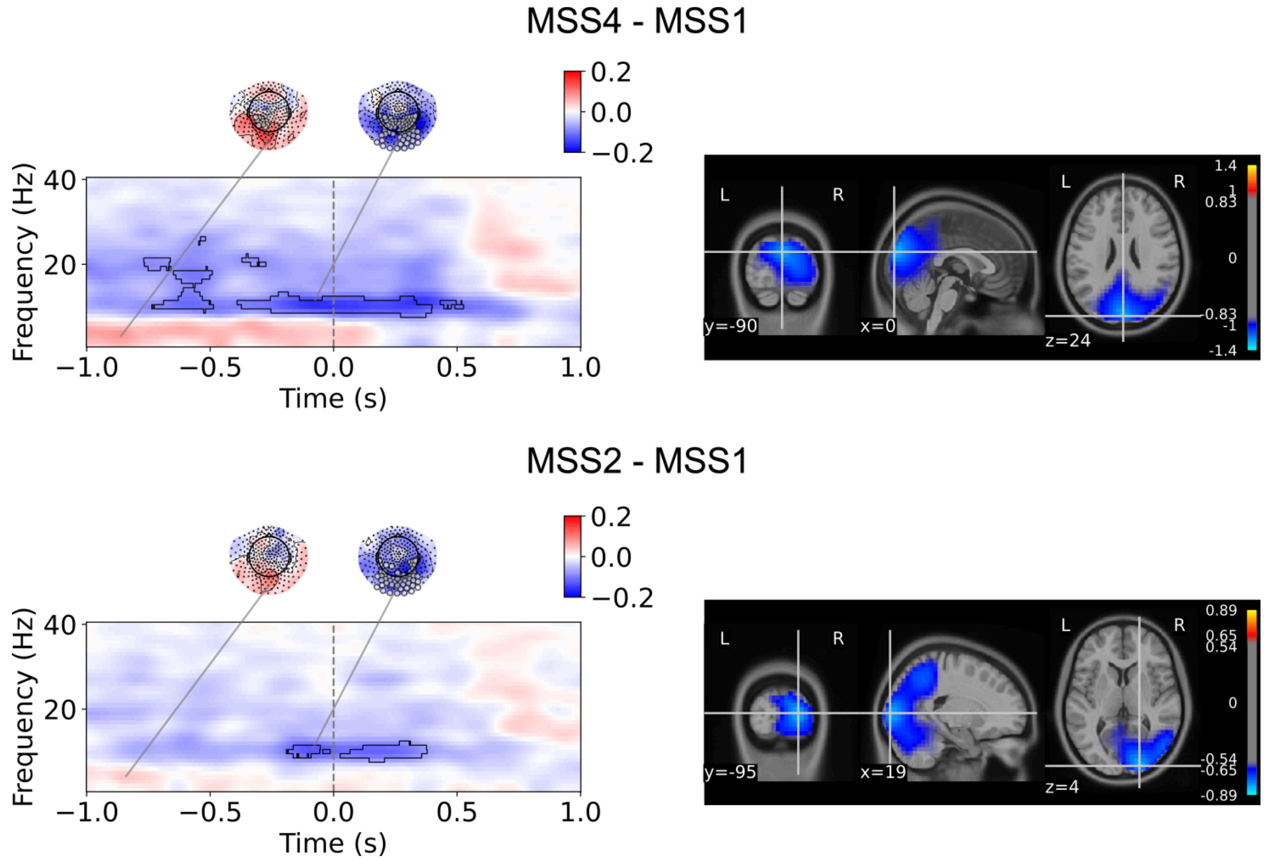


***Figure S1:*** *Difference between the memory load during memorization. Left: temporal frequency response during memory encoding and retention (t=0). Right: Significant power source*

*estimates in the alpha band (8 to 12 Hz) (p = 0.007 for MSS2 - MSS1) (p = 0.002 for MSS4 - MSS1).*

### Figure S2: Encoding power and inter-trial coherence (ITC) on correct vs incorrect trials

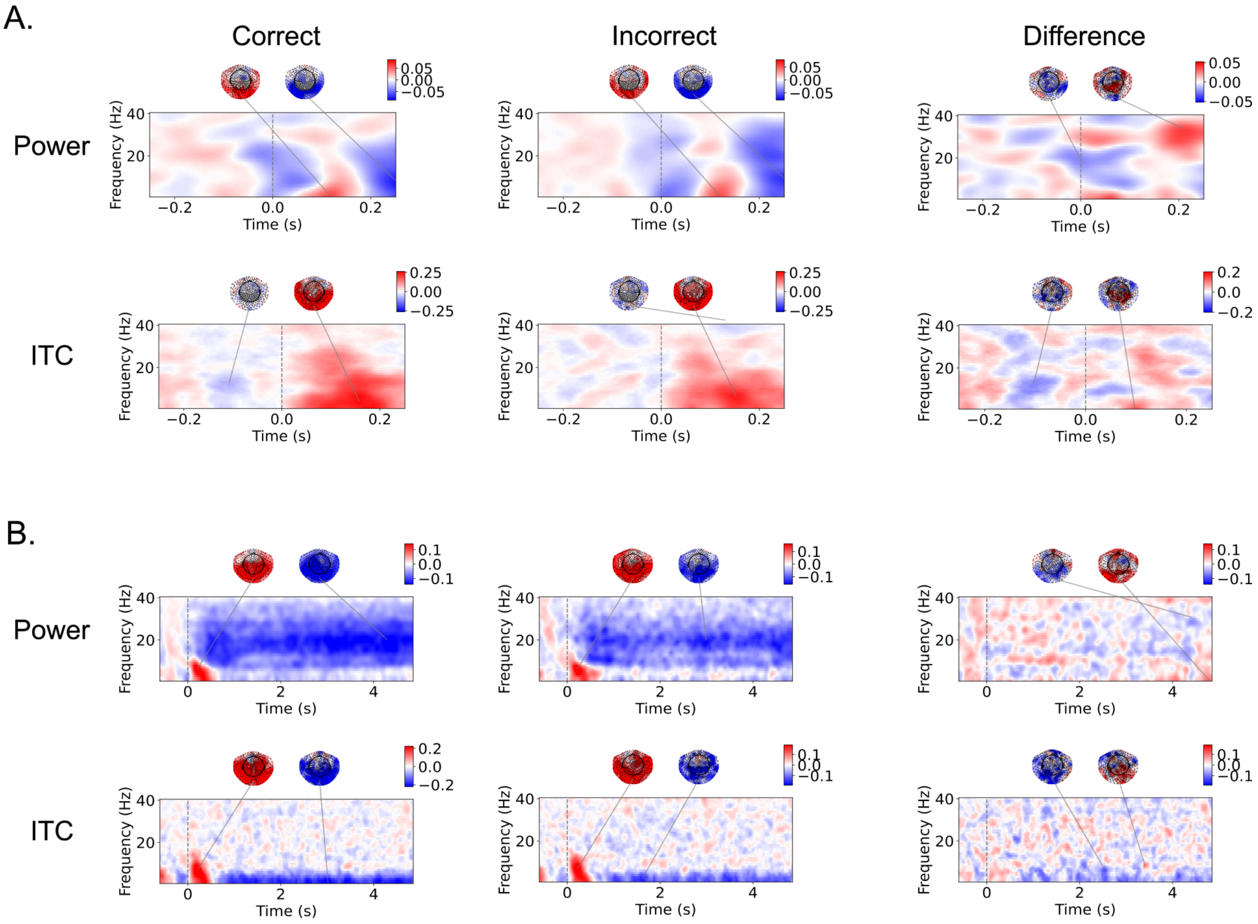


***Figure S2:*** *Power and ITC on encoding screen for MSS 4 trials.* ***A.*** *Results aligned to target and distractor saccades for correct and incorrect responses are shown separately alongside the difference between them. No significant differences were found in the difference between conditions.* ***B.*** *Power and ITC aligned to encoding screen onset for correct and incorrect responses are shown separately alongside the difference between them. No significant differences were found between conditions.*

### Figure S3: Memory load effect on fixations power during visual search

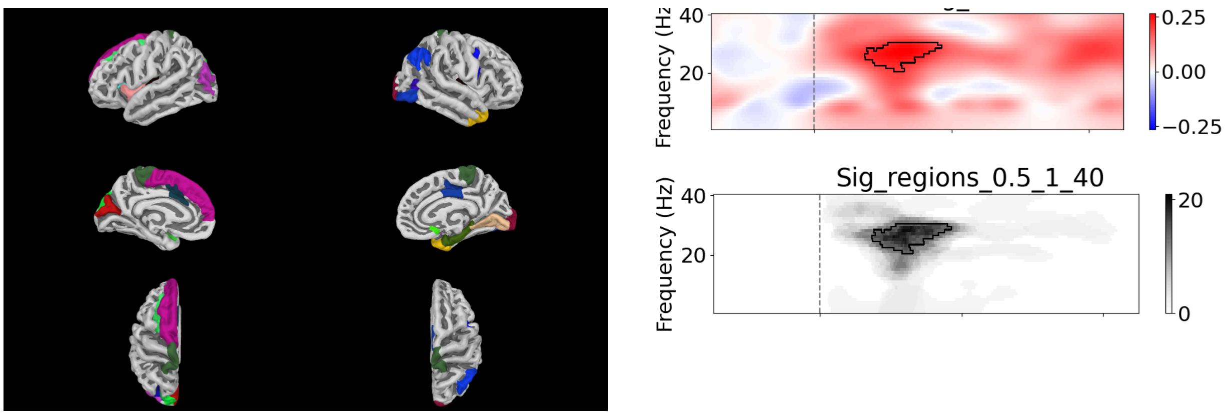


***Figure S3:*** *Working memory load power modulation of saccade response, aligned to saccade onset. Power difference of saccades directed to distractors in visual search of trials under high memory load (MSS=4) and low memory load (MSS=1).*

### Figure S4: Target-related P3m activations in source space

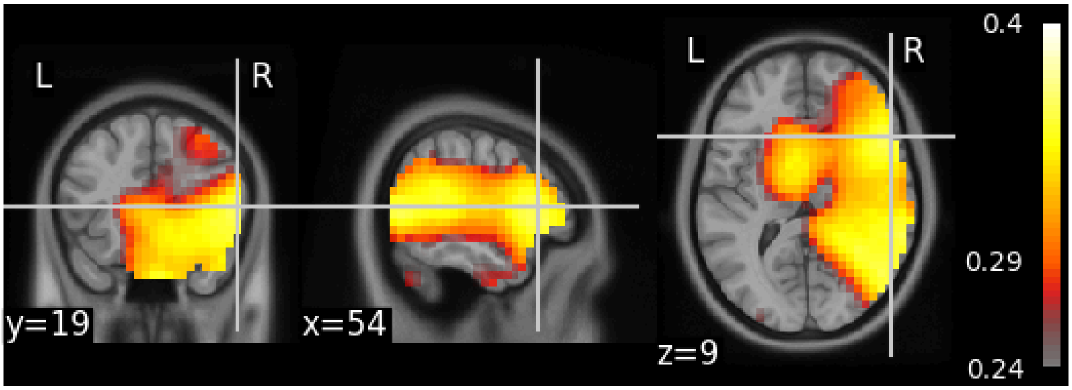


***Figure S4****: Activation map showing the cluster of voxels with significantly different responses to target compared to distractor fixations (p-value = 0.00097). The colorbar indicates the duration on which each voxel presented a significant difference between conditions.*

### Figure S5: Connectivity lateralization analysis

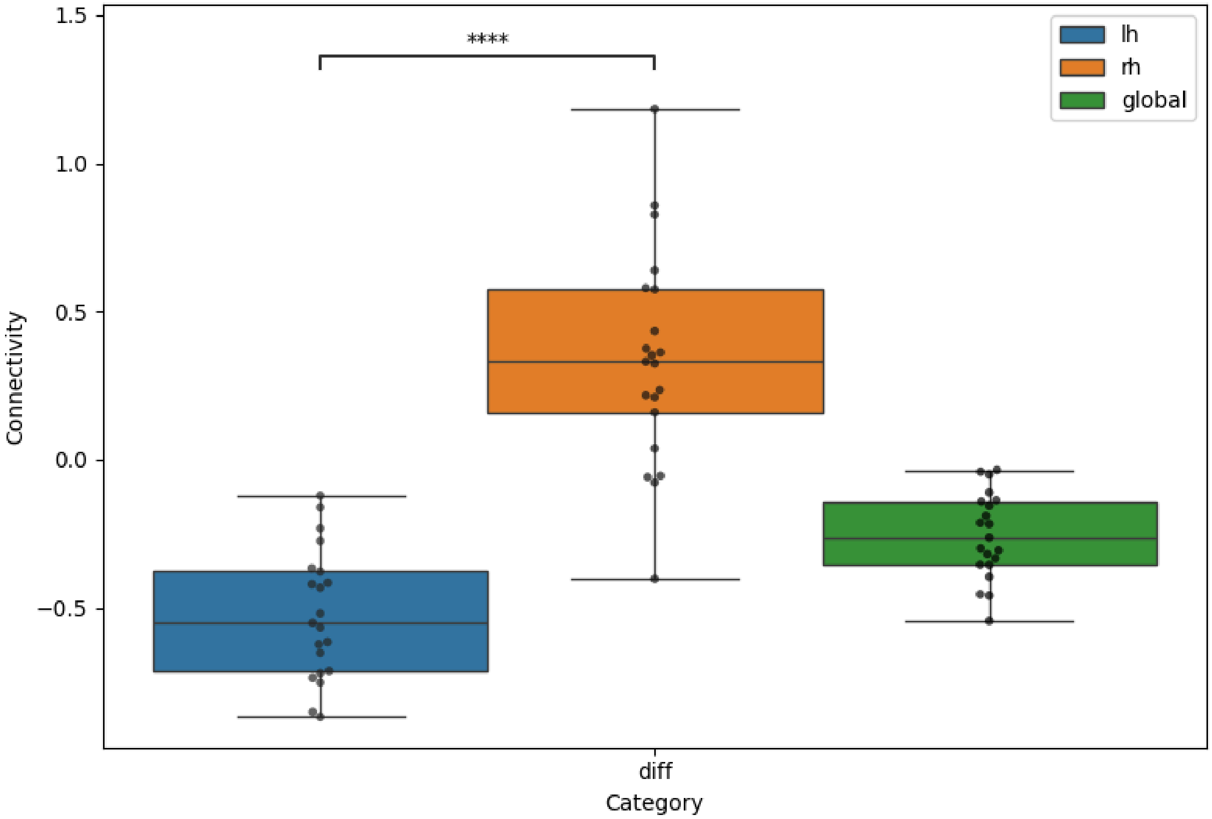


***Figure S5:*** *Wilcoxon signed-rank test results for connectivity lateralization. Each point in the boxplots corresponds to one subject. For each hemisphere, values represent the mean connectivity strength across the 150 strongest links within the left and right hemisphere, respectively. A paired Wilcoxon signed-rank test between left and right hemispheric connectivity indicated a significant difference (W = 0, p = 0.00000095). One-sample Wilcoxon tests were performed against zero for each hemisphere: Both hemispheres were significantly different from zero (W = 0, p = 0.00000095 for left hemisphere), (W = 23, p = 0.0006 for right hemisphere).*
